# Mammalian cell characterisation by non-invasive plate reader assay

**DOI:** 10.1101/2023.03.22.533848

**Authors:** Alice Grob, Chiara Enrico Bena, Roberto Di Blasi, Daniele Pessina, Matthew Sood, Zhou Yunyue, Carla Bosia, Mark Isalan, Francesca Ceroni

## Abstract

Automated and non-invasive mammalian cell analysis is currently lagging behind due to a lack of methods suitable for a variety of cell lines and applications. Here, we report the development of a high throughput non-invasive method for tracking mammalian cell growth and performance based on plate reader measurements. We show the method to be suitable for both suspension and adhesion cell lines, and we demonstrate it can be adopted when cells are grown under different environmental conditions. We establish that the method can inform on effective drug treatment to be used depending on the cell line considered, and that it can support characterisation of engineered mammalian cells over time. This work provides the scientific community with a novel approach to mammalian cell screening, also contributing to the current efforts towards high throughput and automated mammalian cell engineering.

## Introduction

Recent advances in technology and in the automation of cell screening and engineering have made the design and assembly of large libraries of genetic DNA systems possible in mammalian cells^1–3^. This progress builds upon experimental pipelines that have been established to enable the automation of design-build-test analysis of genetic designs in microbes^4,5^. However, the fast identification of desired variants for various applications is only possible if the behaviour of each construct is assessed in the cellular host. It is important to characterise stability, productivity and performance over time, and a key requirement is to measure the impact on cell growth of different constructs.

Whereas high-throughput growth-tracking methods have been developed for bacterial cells^6^, such as simple measurements based on light absorbance^7,8^, automated and non-invasive characterisation of both engineered and non-engineered mammalian cells is lagging behind. Traditional methods for mammalian cell growth characterisation include trypan blue staining, flow-cytometry^9^, automated cell counters^10,11^ and colorimetric assays^12^. However, these methods are disruptive as they rely on a sample of the solution being measured and as such only give a fixed sample measurement in time. Also, they are low throughput and time-consuming, representing a limiting step in the development of high throughput approaches.

Protocols that have been developed for indirect counting of mammalian cells include digital holography^13^, microscopy^14–17^, confluency analysis based on commercially-available plate readers^17^, adoption of magneto sensors^18^ and identification of cell-specific optical density (OD)^19,20^. These are mostly restricted to adherent, monolayered or coated cells, can be expensive and can be limited by the number of samples that one can screen at any one time. The implementation of continuous measurements for both adherent and suspension cells would thus allow for a universal, dynamic, and automated mammalian cell characterisation approach.

To advance the experimental characterisation of mammalian cells, we developed a method for non-invasive, high throughput and automated tracking of mammalian cell growth based on the change in absorbance of the pH indicator phenol red (Figure 1a). This colour indicator is present in a variety of commonly adopted mammalian growth media and is available separately as a dye that can be added to any medium of interest^21^. The acidic and basic forms of phenol red are characterised by absorbance peaks at 560 nm and 430 nm, Abs_560_ and Abs_430_ respectively^22,23^. During mammalian cell growth, phenol red gradually transitions from its basic to its acid form, providing the operator with a quick and visual indication that cells are growing and reaching confluence. Previous work showed that the change in absorbance of phenol red can be used as a valuable method for tracking cell growth in bacteria^24^,^25^ but no one has so far provided a method for mammalian cell tracking using the same principle. Our novel workflow enables characterisation of mammalian cell growth over time, and we demonstrate this working for both suspension and adhesion cell lines, a clear step forward compared to current state of the art. We also confirm that this method can be adopted for a wide range of applications from routine mammalian cell analysis to screening of cell sensitivity to drug treatment, but also for the advancement of high throughput characterisation of engineered constructs in these cells, thus making it relevant for both basic and applied research.

**Figure 1.**
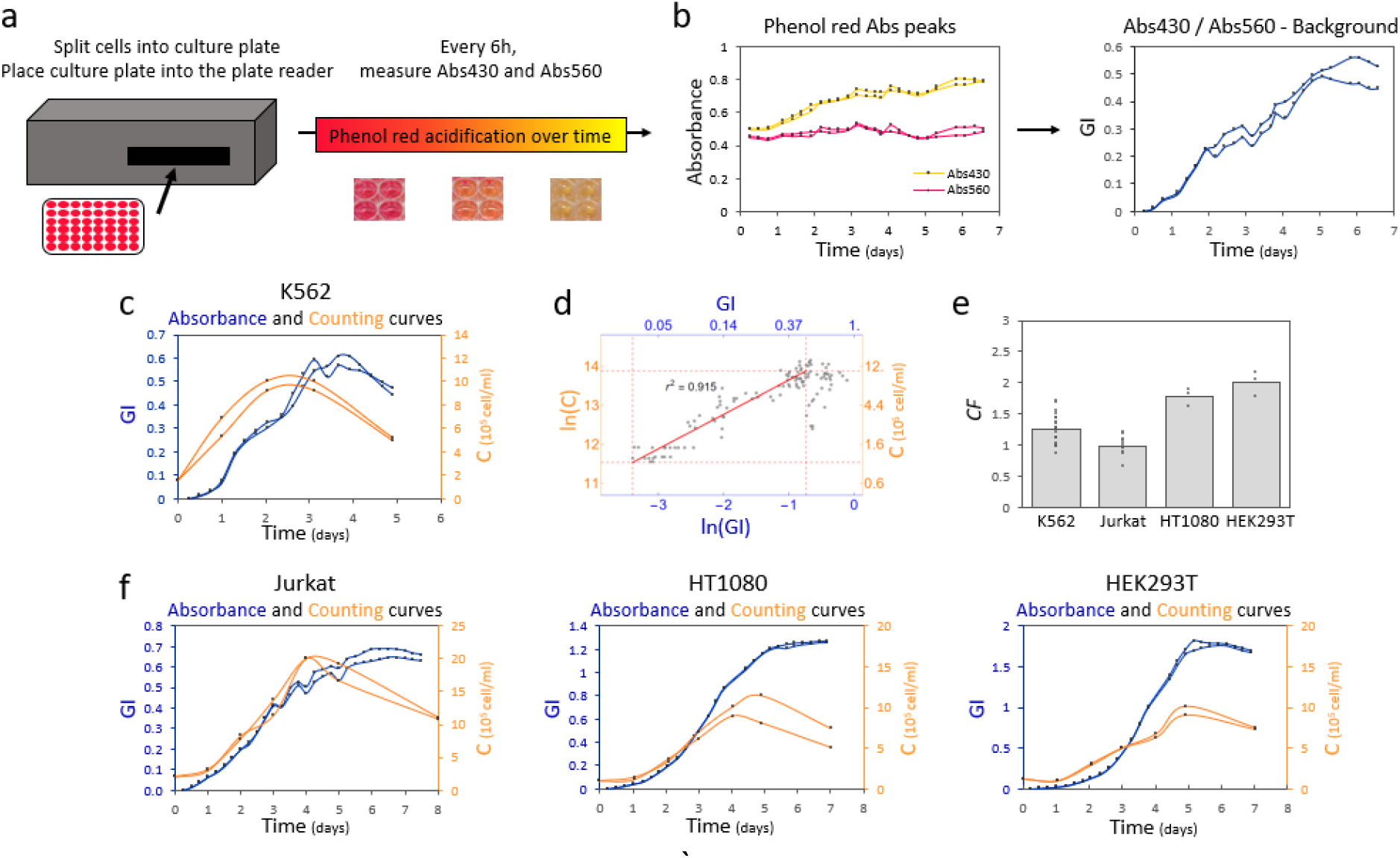
A plate-reader assay based on the absorbance of the pH indicator phenol red can be used to dynamically characterise the growth of mammalian cell lines. **a)** Schematic of phenol red-based mammalian growth assay. **b)** Representative output of the assay with dynamic of Abs_430_ (yellow) and Abs_560_ (pink) (left). The resulting ratio of Abs_430_ over Abs_560_ normalised to the background yields the GI profile over time for K562 cells (right). **c)** Representative growth curves resulting from phenol red acidification (GI, blue, left vertical axis) and cell counts (orange, right vertical axis) of K562 cells. All replicates can be found in Related File 1. **d)** Correlation between K562 cells GI and C. The red line is the best fit of data within the linear region, i.e. ln(C) = 14.5 + 0.88*ln(GI) with a coefficient of determination r^2^=0.915. Red dashed lines represent the edges of the linear region, where the two plotted variables scale linearly. **e)** Bar plot of *CF* for suspension cells (K562 and Jurkat) and adherent cells (HT1080 and HEK293T). The height of the bar represents the mean value of the single replicates shown as black dots. **f)** Representative profiles of GI (blue, left vertical axis) and C (orange, right vertical axis) for Jurkat, HT1080 and HEK293T cells. All replicates can be found in Related File 1. Numbers of biological repeats for each sample are reported in Table S3. Data analysis is described in the Methods section and in Supplementary Note 1.

### A phenol red-based growth assay enables plate reader tracking of mammalian cell growth

We started by considering the suspension cell line K562^26^ and designed a routine protocol where cells are first seeded at their recommended seeding density in a 48-well plate. The plate is then inserted into a plate reader where temperature is set at 37°C and CO_2_ at 5%. Absorbance readings for the basic (Abs_560_) and acidic (Abs_430_) forms of phenol red are performed every six hours for up to nine days (Figure 1a).

It was previously suggested for microbes, that a growth index (GI) can be identified by the ratio between Abs_430_ and Abs_560_^24,25^, which yields a characteristic sigmoidal growth profile (Figure 1b, Figure S1a-c). To confirm that a GI is also a reliable proxy of cellular growth in mammalian cells, we performed parallel cell counts every 24 hours for each sample, using a standard automated cell counter (see Methods section). We reasoned that this would allow us to compare the growth profiles yielded by the two methods and to benchmark against one of the techniques most adopted for following mammalian cell growth. As expected for batch cell cultures, cell concentration (C) yielded a typical sigmoidal curve, inclusive of lag, exponential and stationary phases of growth (Figure 1c). Plate reader growth curves showed an analogous growth profile (Figure 1c, Related File 1).

Specifically, we identified a range for which the logarithm of C (ln(C)) and the logarithm of GI (ln(GI)) follow a linear relation and that this corresponds to the exponential phase of growth (Figure 1d, red line, Table S1 and Supplementary Note 1). Thanks to this, it was possible to compute the growth rate from plate reader (μ_p_) and cell counts (μ_c_) in the same way, i.e. by computing the slope of the data within the exponential phase as a function of time (see Supplementary Note 1 for data analysis details). In order to establish the relationship between μ_p_ and μ_c_, we developed a protocol for automated analysis that enabled the calculation of the growth rate from each dataset (see Supplementary Note 1 and Related File 2). We named conversion factor (*CF*) the ratio between μ_p_ and μ_c_.

We found that *CF* for K562 cells is around 1, suggesting that the indirect μ_p_ computed through GI is a proxy for the effective μ_c_ of the population obtained by direct cell counting (Figure 1e). This relation is valid within the exponential phase, when μ_c_ and μ_p_ scale linearly to each other, as shown in Figure 1d. Analogously to the relation observed in bacteria between OD and cell concentration^8^, linearity between the two quantities is lost when cells start saturating. As evident from Figure 1c, after saturation, the growth curve for C decreases more rapidly than the one for GI. Additionally, the slope of the linear fit between ln(GI) and ln(C) provides information on the relation between the actual number of cells and the detected absorbance ratio (Table S2). For K562 cells, a slope value close to 1 (i.e. 0.88) suggests that GI scales almost linearly with C. Thus, for K562 cells growing in standard conditions, GI in exponential phase can be used as a proxy for estimating C (see Supplementary Note 1).

### Suspension and adhesion cell lines can be characterised by plate reader-based analysis

We next set out to verify whether it was possible to apply this method to a range of cell lines, and specifically to both suspension and adherent cells. We thus adapted our protocol to characterise the growth of a second suspension cell line (Jurkat^27^) and two cell lines growing in adhesion (HT1080^28^ and HEK293T^29^). As shown in Figure 1f, when the cells are grown in standard growth medium at 37°C, the GI captures the trend of the growth profiles of all of these cell types reliably (Figure 1f, Figure S1d-f, Figure S2), again leading to a linear relation between ln(GI) and ln(C), within the exponential phase of growth (Figure S3 and Table S1).

When we compared the *CF* for the four different cell lines, we noticed that the *CF* for Jurkat cells is also close to 1, as for K562 cells, while both adhesion cell lines show an average CF closer to 2 (Figure 1e, Table S4 and Supplementary Note 1). To explain this difference, we considered that both K562 and Jurkat cells are cultured in RPMI medium (containing 5mg/ml phenol red), while HT1080 and HEK293T cells are cultured in DMEM (containing 15mg/ml phenol red). Indeed, we noticed that Abs_560_ follows a very different trend for cells grown in DMEM (Figure S2a, S2b), compared to RPMI (Figure S1a, S1b), with a more pronounced decreased over time in the former case. This is then reflected in a wider GI variation (Figure S2e and S2f). Experimental evidence previously showed that DMEM, and specifically Abs_560_ in this medium, is more sensitive to pH change if compared to RPMI, supporting our hypothesis that the difference in *CF* is directly linked to a media-dependent effect that leads to a different profile in Abs_560_^30^.

Despite this, the linear relation between ln(GI) and ln(C) observed for both suspension cell lines still holds true for adhesion cells when the exponential phase of growth is considered, as shown in Figure S3. Thus, a *CF* can be experimentally calculated for any cell line of interest where the linearity between ln(GI) and ln(C) is conserved. Overall, these results, suggest that μ_p_ can be treated and considered as a direct proxy for the actual cellular μ_c_.

### Phenol red-based growth assay enables mammalian growth tracking under different conditions

Once we established that our method could reliably follow the growth profiles of different cell types, we investigated if it could be adopted for characterisation of cell growth across different culture conditions. Temperature and carbon source are two key parameters in mammalian process optimisation, and they have been previously shown to impact bioproduction and therapeutic applications^31–33^. Proving that our protocol is robust to varying environmental conditions is thus an essential requirement to confirm it can be applied to wider mammalian cell analysis.

We focused on comparing K562 and Jurkat cells growing at 37°C and 33°C, in the presence of glucose (Glu) or mannose (Man) (Figure 2 and Figure S4). GI profiles still reliably captured cell count profiles, for both cell lines growing at 33°C (Figure 2a and S4a) and when Man was used as carbon source (Figure 2b, 2c, S4b and S4c). Data analysis by our automated pipeline confirmed that changing growth conditions did not affect the linear relation previously observed between ln(GI) and ln(C) (Figure 2d-f and S4d-f). Importantly, the average value of *CF* for both cell lines did not change (Figure S5a, Table S5 and Related File 2).

**Figure 2.**
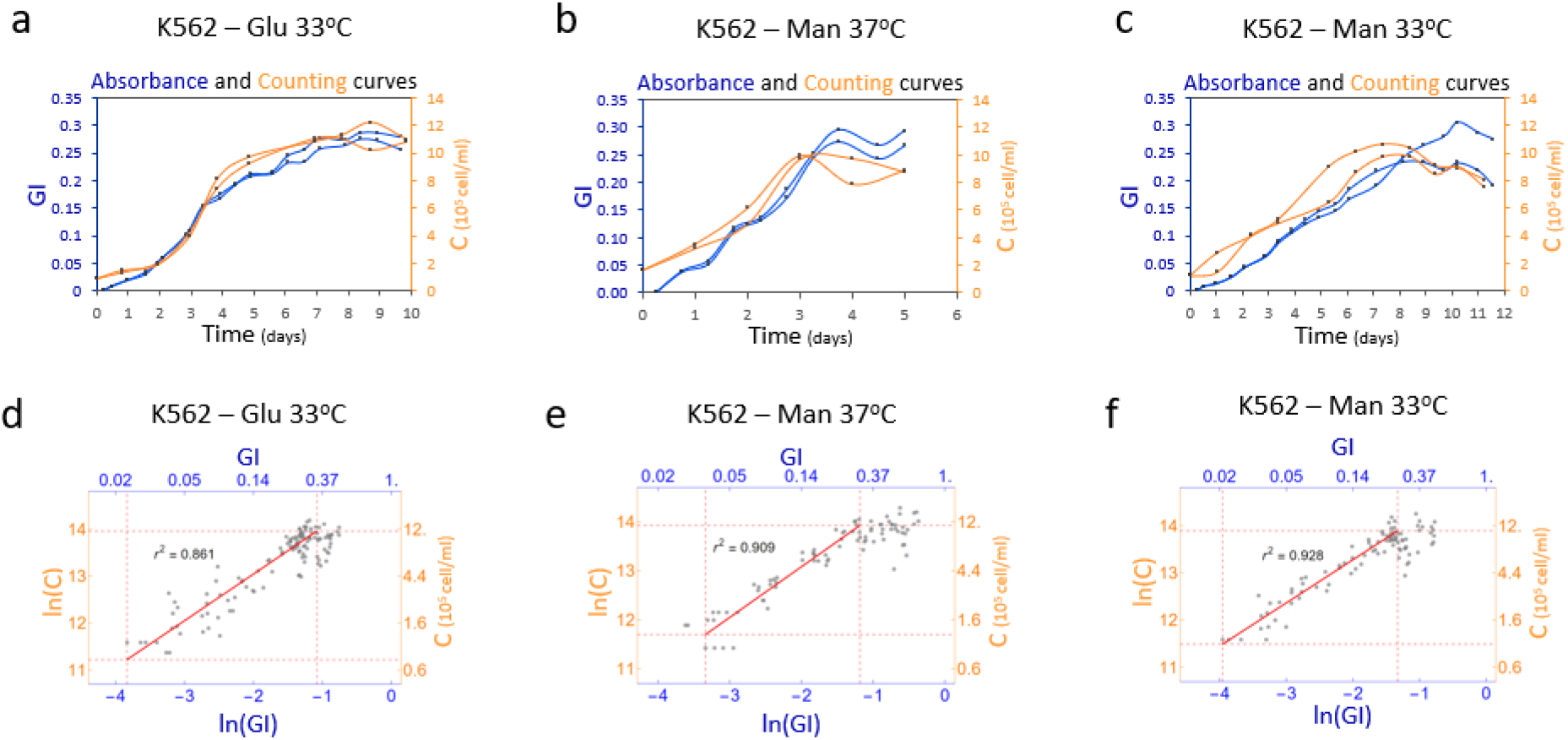
Growth of mammalian cell lines under different conditions can be characterised by a plate reader assay. Representative GI (blue, left vertical axis) and C (orange, right vertical axis) profiles over time of K562 cells grown with Glu **a)** at 33°C or Man at either **b)** 37°C or **c)** 33°C. All replicates can be found in Related File 1. Correlation plot for ln(GI) and ln(C) for K562 cells grown with Glu **d)** at 33°C or Man at either **e)** 37°C or **f)** 33°C, indicating r^2^ values of 0.861,0.909 and 0.928 in exponential phase, respectively. Red dashed lines represent the edges of the linear window, while the red solid line correspond to the linear fit. Fit equations are reported in Table S2. Numbers of biological repeats for each sample are reported in Table S3. Data analysis is described in the Methods section and in Supplementary Note 1.

Based on these results, we decided to verify how automatable our pipeline is and verify if, once we established GI, μ_p_ and *CF* for a given cell line and condition, it could be possible to infer μ_c_ for subsequent experiments where only plate reader measures of GI are performed in the same conditions. We thus used the calculated μ_p_ and *CF* from an initial set of measures for K562 and Jurkat cells growing at 37°C and 33°C, in the presence of Glu or Man, to estimate μ_c_ for subsequent plate reader runs and compared the estimated values (μ_cp_) with the actual μ_c_ calculated by parallel cell counting (Figure S5b). μ_cp_ closely matched the measured μ_c_, thus confirming that once an initial characterisation is performed, the identified *CF* can be adopted to infer information on actual μ_c_ of the cells in further experiments without need for cell counting.

### High throughput screening of therapeutic drugs enables identification of cell linespecific response

Doxorubicin (doxo) is a well-known chemotherapeutic agent adopted in anticancer medications to treat, amongst others, breast cancer and leukaemia^34^. Doxo inhibits cell growth via interference with DNA replication and RNA synthesis, leading to cell death^35^. While traditional methods for identification of effective doxo concentrations, and for screening sensitivities of different cell lines to the drug, involve manual or automated cell counting, we reasoned that our platform could provide an improved workflow for testing.

Here, we applied our protocol to test the effect of doxo on two of the four cell lines considered, namely HT1080 (known to be doxo sensitive^36,37^), and K562, selected as test-bed of a leukaemia cell line^26^. Our data confirmed that HT1080 cells are indeed sensitive to doxo, as demonstrated by the continuous change in GI profile and corresponding stepwise decrease in μ_p_ when increasing doxo concentrations are added (Figure 3a and 3b top). Specifically, we observed that μ_p_ for HT1080 cells decreases to ~80% and further to ~40% when 25nM and 500nM doxo are adopted, respectively, compared to the untreated control (Figure 3b). By contrast, K562 cells displayed less sensitivity to doxo treatment, displaying a constant GI and μ_p_ (Figure 3a and 3b bottom) with cells still displaying 90 to 100% μ_p_, in the presence of doxo concentrations up to 500nM (Figure 3b). Effective decreases in cell growth were observed when doxo was increased to 750nM, and then to 1μM, where cells displayed ~76% and ~65% viability compared to the control, respectively. The method effectively supported characterisation of the cell line-specific response to doxo treatment and informed on a drug concentration range for effective cell growth inhibition.

**Figure 3.**
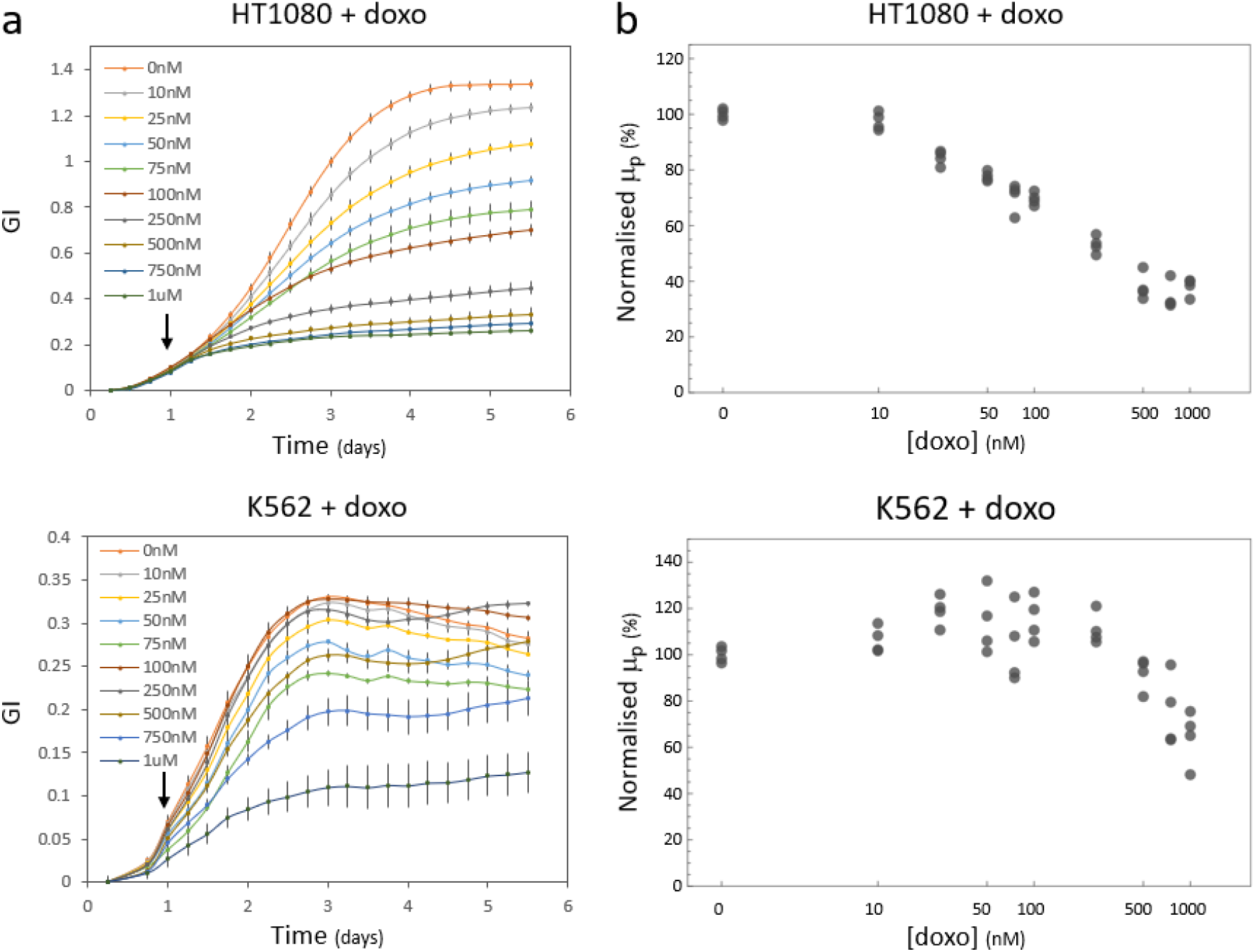
Growth of mammalian cell lines treated with doxorubicin can be characterised by a plate reader assay. **a)** Response of HT1080 and K562 cells to treatment with increasing doxo concentrations (0nM to 1μM). Doxo was added after 24 hours from the start of the plate reader assay (arrow). **b)** Relative μ_p_ values of HT1080 and K562 cells treated with 0-1000nM doxo (axis in log scale) were normalised to the average value of μ_p_ for cells treated with 0nM doxo. Number of biological repeats for each sample are reported in Table S3. Data analysis is described in the methods section and in Supplementary Note 1.

### Automated tracking of engineered mammalian cells and single cell construct performance

Finally, we thought to apply our method to the characterisation of engineered mammalian cells, considering that high throughput and automated analysis of engineered hosts is much needed within the biotechnology and synthetic biology communities. To provide a proof-of-concept of the suitability of our method for this task, HEK293T-LP cells, a HEK293T cell line bearing a landing pad (LP) cassette integrated in the AAVS1 locus (available from *Matreyek et al*^38^) were selected. The LP cassette codes for a BFP expressed under an inducible Tet promoter. The LP also bears a second transcriptional unit expressing the rtTA transactivator under the control of a constitutive EF1α promoter (Figure 4a). When doxycycline (dox) is present, the rtTA transcription factor activates the Tet promoter and BFP is expressed. Conversely, when dox is absent, the cassette is off. Since the switch is known to be sensitive to different dox levels, we reasoned it could work as a good proxy for genomic constructs with different expression strengths, thus providing a good test bed for our method.

**Figure 4.**
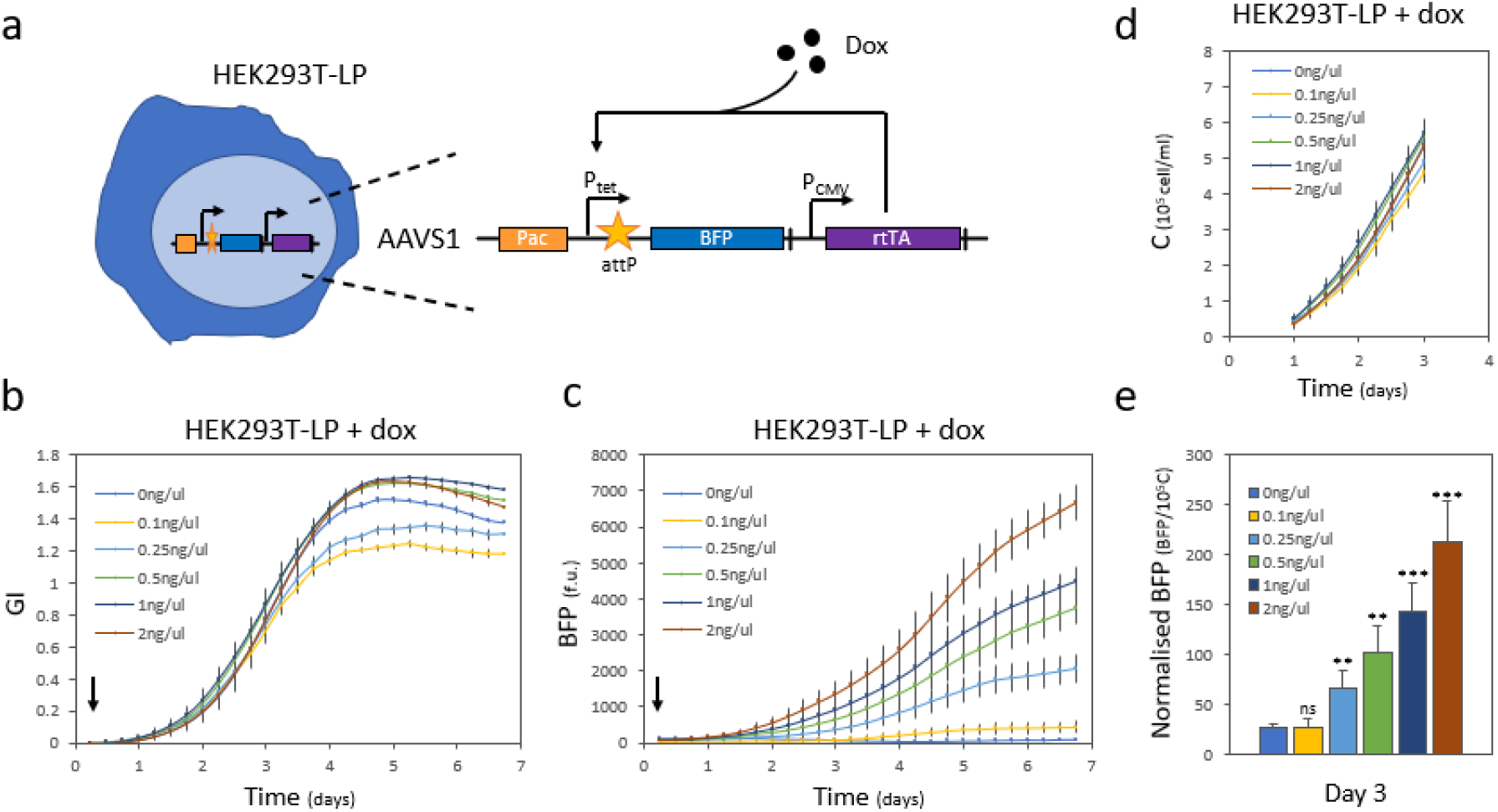
Gene expression from a landing pad cassette integrated in HEK293T cells can be characterised over time with a plate reader assay. **a)** Schematic of the landing pad in HEK293T-LP cell line from *Matreyek et al^38^*. A cassette expressing the dox-responsive rtTA transactivator and the inducible Ptet-BFP transcriptional unit is integrated in the AAVS1 locus of HEK293T cells. **b)** GI and **c)** BFP fluorescence of HEK293-LP cells can be followed over a one-week time frame for samples induced with increasing concentrations of dox (0ng/μl to 2ng/μl). Addition of dox at T0 is indicated by an arrow. **d)** Estimated C, derived from GI, within the linear time frame between the two variables (i.e. one to three days). **e)** Fluorescence per cell can be calculated by normalising the total BFP fluorescence on the estimated C at three days post induction. A two-sided t-test indicated that normalised BFP levels increase significantly in cells treated with 0.25ng/μl dox and above (ns, non-significant: p>0.5, **: p<0.01 and ***: p<0.001). Numbers of biological repeats for each sample are reported in Table S3. Data analysis is described in the Methods section and in Supplementary Note 1.

We initially characterised the cell line following our protocol as outlined in Figure 1a (Figure S6a-c). Performing plate reader measurements and cell counts in parallel (Figure S6d), we confirmed linearity between ln(GI) and ln(C) for this additional cell line (Figure S6e) and established its specific *CF* (Figure S6f). We then characterised the response of the LP inducible cassette to dox induction. To do so, the HEK293T-LP cells were seeded in a 48-well plate and were allowed to grow in the plate reader for one week after induction with different concentrations of dox. GI for the different samples and total BFP fluorescence were captured (Figure 4b and 4c). As expected, increased dox induction led to increasing BFP levels.

Bacterial OD is routinely used to infer normalised protein expression per cell. This is an essential practice to compare samples and conditions, as it allows to take into account differences in growth and in cell number when characterisation of a construct functionality is sought. Given that ln(GI) is linear with ln(C) during exponential growth, we thought to exploit such relation to calculate the C value corresponding to a given GI value (Figure 4d), similarly to what we did for μ_c_ estimations previously (see Supplementary Note 1 for details). This enabled us to normalise the protein expression on the estimated number of cells at a given moment in time (i.e. three days), within the linearity window, and obtain an estimated quantitative measure of expression levels per cell (Figure 4e).

In the work published by *Matreyek et al*, authors performed flow cytometry measurements and characterisation of the induction dynamic of the HEK293-LP cell line, looking at BFP signal expression over time. They monitored that cells induced with 2ng/μl dox display an increase in BFP expression per cell up to five days. Our normalised BFP data (Figure S7) show a similar increase in BFP per cell over time after induction. In conclusion, our method led us to similar single cell construct characterisation but via a non-invasive and high throughput method that enabled both growth and fluorescence tracking with no requirement for manual handling.

## Discussion

The ability to track mammalian cell growth and performance over time, adopting automated and non-invasive protocols, is key to supporting current advances in mammalian cell research. In this study, we developed a first-of-its kind plate-based high throughput method for the characterisation of mammalian cell growth. The method is based on the change in absorbance observed for the colour indicator phenol red during growth in mammalian cultures. It consists of four main steps, in which (i) cells are first seeded with a desired concentration and growth medium into a 48-well plate that is then inserted into a plate reader with temperature and CO_2_ control; (ii) measurements of Abs_430_ and Abs_560_ are performed every six hours for one week; (iii) a ratio of these absorbances is computed in order to generate a GI curve; (iv) automated data analysis is used to identify the growth rate of the GI curve in exponential phase. An initial calibration is required for any new cell line to be analysed, and this can be done by performing cell counts every 24h in parallel to the initial plate reader measurements. Then the automated data analysis workflow provided here supports analysis of the cell counts and plate reader curves to identify their corresponding growth rates. The resulting ratio between the calculated growth rate in a plate reader, μ_p_, and the growth rate from cell counts, μ_c_, defines a conversion factor, *CF*, which holds in exponential phase, i.e. when ln(GI) and ln(C) scale linearly.

We started here by showing that our method enables tracking of both suspension and adhesion cell lines, going beyond existing methods currently restricted to one of the two cell types^15,17^. We then showed that together with standard growing conditions, the method can be used to characterise cell growth and performance when temperature and medium composition are changed. The method also successfully provides characterisation of cell-specific responses to chemotherapeutic drug treatment and identification of effective working drug concentrations. Finally, we demonstrated that engineered mammalian cells can be characterised in their growth and fluorescent outputs when an integrated cassette is expressed. Importantly, we provide evidence that, once *CF* is established for a given cell line in a specific condition, the method can be adopted to infer actual μ_c_ and cell numbers from plate reader measurements alone. By adopting *CF* to convert plate reader data into cell counts, the protocol enables quantification of construct and cell performance over time when different induction is sought. This strengthens the relevance of the methods widening its potential applications in mammalian cell analysis. Our method has the potential to greatly support advancements in the automation of mammalian cell screening both in basic and applied research. For the latter, while for several micro-organisms it is now possible to perform automated library assembly and parallel testing of high number of variants^39–41^, the uptake of the same workflow in mammalian cells is lagging behind. Design and assembly of large construct libraries is now possible for mammalian systems thanks to the development of toolkits that support modular and high throughput construct generation^42,43^, but the screening of such variants within their target host is slowed down by the low throughput protocols that were previously available. Biofoundries have been created over the last decade to address this need, specifically aiming to introduce automation into the well-known design-build-test cycle of synthetic biology^44^. In parallel, important technological advances have arisen, like the ones showcased by companies such as Berkley lights that allow automated cell line development and clone selection^45,46^. However, such technologies are still currently inaccessible to lab-end users, due to their high costs and need for dedicated lab space.

Methods which are more easily accessible, such as the one developed here, will thus be pivotal in supporting advancement of mammalian cell analysis, and in enabling a more dynamic, automated and higher throughput characterisation of mammalian cells than was thus far possible.

## Material and Methods

### Cell culture

Suspension cells K562 (ATCC CCL-243) and Jurkat (ATCC TIB-152), and adherent cells HT1080 (ATCC CCL-121) and HEK 293T AAVS1 LP (HEK293T-LP, kind gift from D.M. Fowler) were cultured respectively in RPMI medium 1640 (Gibco) or DMEM medium (Gibco) with 10% foetal bovine serum (FBS) (Gibco). For characterisation of growth in different conditions, suspension and adherent cells were pre-cultured for two weeks in their respective glucose-free medium with 10% FBS and 25mM Man (Sigma). This concentration of Man was also adopted for plate base and cell count measurements.

### Plate reader assay

Suspension and adherent cells were resuspended in fresh medium at 10^5^ cells/ml and plated into a 48-well plate (Greiner). To limit evaporation, phosphate buffered saline (PBS; Sigma-Aldrich) was added in-between wells and the plate was sealed with parafilm on its long sides, leaving the short sides for CO_2_ exchange. The plate was then placed in a SPARK or an Infinite M200 Pro microplate reader (Tecan) and incubated at 33°C or 37°C ± 0.5 with 5% ± 0.5 CO_2_ for up to 15 days. Every six hours, the plate of suspension cells was orbitally shaken for 20 seconds, incubated for 5 seconds and then Abs_430_ and Abs_560_ were measured; while for adherent cells Abs_430_ and Abs_560_ were measured directly every six hours. **Cell counting.** Cell counting was performed in parallel to the plate reader assay. Every 24h, the 48 well plate was taken out of the microplate reader, 5μl of suspension cells were stained with 0.2% trypan blue (Gibco) and counted using a TC20^TM^ automated cell counter (Bio-Rad). Adherent cells were detached using 0.05% Trypsin-EDTA (Gibco) then inactivated by v/v DMEM medium. Cell counting of these cells was performed using a Nucleocounter NC-250™ (Chemometec). Remaining cells were removed from the well and replaced by PBS to minimise evaporation. Subsequently, the plate was sealed with parafilm again on its long sides and replaced inside the microplate reader prior to restarting the plate reader assay.

### Growth curves

For suspension cell lines, cell counting, and absorbance measurements were performed from the same well, from seeding to saturation. Thus, the growth curves obtained for these cells correspond to measurements over time from a single well. On the contrary, for adherent cells, each time point corresponds to a different well due to the need for trypsinization and detachment of the cells before counting. For both C and GI growth curves (Figure 1f, S6d) each time point thus corresponds to the measurement of a given well at the given time. In parallel to the wells used for cell counting, two wells were measured by sole plate reader for the entire duration of the experiment. GI values were normalised to their initial values at six hours, GI_0_ (see Supplementary Note 1).

### Linear correlation between GI and C

To infer the relation between GI and C, we analysed ln(C) as a function of its correspondent ln(GI) values. All the values of independent growth curves belonging to the same experimental condition (i.e. cell line, supplied sugar and temperature) were pulled together and analysed in two main steps. First, the range of linearity between the two variables was determined in an automatic and unbiased way; second, the data belonging to this linear range were fitted with a linear fit, obtaining the relation between ln(GI) and ln(C). All data were analysed through custom made scripts in *Wolfram Mathematica*. The code used for data analysis, description of it and source data file are available as Related File 3. See Supplementary Note 1 for further details.

### Conversion factors

The linearity between ln(GI) and ln(C) allows to convert the exponential growth rate of cells growing in the plate reader, μ_p_, into that obtained through cell counting, μ_c_, with a proportion. The conversion factors that allow this are computed as *CF* = μ_p_/μ_c_. See Supplementary Note 1 for the computation of the exponential growth rate.

### Doxorubicin assay

HT1080 and K562 cells were plated into a 48 well plate at 10^5^ cells/ml. The plate reader assay was started and let run for 24h. The plate was then removed from the microplate reader and 0 to 1 μM doxo (Fisher Scientific) was added to the plate. The plate was then placed back into the plate reader and absorbance measures performed for six more days.

### BFP expression assay

HEK293T-LP cells sourced from *Matreyek et al^38^* were plated into a 48 well plate according to the plate reader assay and treated with 0 to 2 ng/μl dox (Sigma) to induce BFP expression. The plate was then placed into the microplate reader and the assay started. Every six hour, following Abs_430_ and Abs_560_ measurements, BFP fluorescence was also recorded (λ_ex_=381, λ_em_=445) for seven days.

### Statistical analysis

A two-sided t-test was used to assess the statistical significance of *CF* resulting from different growing conditions, μ_cp_, and BFP expression levels.

## Author contribution

AG, MI and FC conceptualised the research. AG, RDB, DP, MS, ZY performed the measurements. CEB and CB performed mathematical and data analysis. FC, MI and CB contributed funding. AG, CEB, MI and FC wrote the manuscript. All authors read and edited the manuscript.

## Acknowledgements

This work was supported by the New Investigator Award no. WT102944 from the Wellcome Trust U.K. (WT102944 to A.G. and M.I.) and the Biotechnology and Biological Sciences Research Council (grant BB/V00882X/1 to A.G and F.C.). FC, CEB and CB acknowledge the support of the Royal Society International Exchanges 2018 Round 1 grant IES \R1\180027. The authors would like to thank *Matreyek et al* for providing the HEK293T-LP cell line adopted in this work, Paul Freemont and Jose Jimenez for feedback on the manuscript.

## References

1 James, J. S. et al. Automation and Expansion of EMMA Assembly for Fast-Tracking Mammalian System Engineering. Acs Synth Biol 11, 587–595, doi:10.1021/acssynbio.1c00330 (2022).

2 Kramme, C. et al. An integrated pipeline for mammalian genetic screening. Cell Rep Methods 1, 100082, doi:10.1016/j.crmeth.2021.100082 (2021).

3 Di Blasi, R., Zouein, A., Ellis, T. & Ceroni, F. Genetic Toolkits to Design and Build Mammalian Synthetic Systems. Trends Biotechnol, doi:10.1016/j.tibtech.2020.12.007 (2021).

4 Chao, R., Mishra, S., Si, T. & Zhao, H. Engineering biological systems using automated biofoundries. Metab Eng 42, 98–108, doi:10.1016/j.ymben.2017.06.003 (2017).

5 Hillson, N. et al. Building a global alliance of biofoundries. Nat Commun 10, 2040, doi:10.1038/s41467-019-10079-2 (2019).

6 Kurokawa, M. & Ying, B. W. Precise, High-throughput Analysis of Bacterial Growth. J Vis Exp, doi:10.3791/56197 (2017).

7 Hall, B. G., Acar, H., Nandipati, A. & Barlow, M. Growth rates made easy. Mol Biol Evol 31, 232–238, doi:10.1093/molbev/mst187 (2014).

8 Stevenson, K., McVey, A. F., Clark, I. B. N., Swain, P. S. & Pilizota, T. General calibration of microbial growth in microplate readers. Sci Rep 6, 38828, doi:10.1038/srep38828 (2016).

9 Kumar, N. & Borth, N. Flow-cytometry and cell sorting: an efficient approach to investigate productivity and cell physiology in mammalian cell factories. Methods 56, 366–374, doi:10.1016/j.ymeth.2012.03.004 (2012).

10 Dittami, G. M., Sethi, M., Rabbitt, R. D. & Ayliffe, H. E. Determination of mammalian cell counts, cell size and cell health using the Moxi Z mini automated cell counter. J Vis Exp, doi:10.3791/3842 (2012).

11 Carvell, J. P., Thomson, K. M. A new automated cell counter for mammalian cell culture assessment. BMC Proc 9, doi:https://doi.org/10.1186/1753-6561-9-S9-P51 (2015).

12 Vega-Avila, E. & Pugsley, M. K. An overview of colorimetric assay methods used to assess survival or proliferation of mammalian cells. Proc West Pharmacol Soc 54, 10–14 (2011).

13 Molder, A. et al. Non-invasive, label-free cell counting and quantitative analysis of adherent cells using digital holography. J Microsc 232, 240–247, doi:10.1111/j.1365-2818.2008.02095.x (2008).

14 Fracassi, C., Postiglione, L., Fiore, G. & di Bernardo, D. Automatic Control of Gene Expression in Mammalian Cells. Acs Synth Biol 5, 296–302, doi:10.1021/acssynbio.5b00141 (2016).

15 Busschots, S., O’Toole, S., O’Leary, J. J. & Stordal, B. Non-invasive and non-destructive measurements of confluence in cultured adherent cell lines. MethodsX 2, 8–13, doi:10.1016/j.mex.2014.11.002 (2015).

16 Odeleye, A. O. O., Castillo-Avila, S., Boon, M., Martin, H. & Coopman, K. Development of an optical system for the non-invasive tracking of stem cell growth on microcarriers. Biotechnol Bioeng 114, 2032–2042, doi:10.1002/bit.26328 (2017).

17 Lanigan, T. M. et al. Real time visualization of cancer cell death, survival and proliferation using fluorochrome-transfected cells in an IncuCyte((R)) imaging system. J Biol Methods 7, e133, doi:10.14440/jbm.2020.323 (2020).

18 Shekhar, S., Karipott, S. S., Guldberg, R. E. & Ong, K. G. Magnetoelastic sensors for real-time tracking of cell growth. Biotechnol Bioeng 118, 2380–2385, doi:10.1002/bit.27680 (2021).

19 Aijaz, A., Trawinski, D., McKirgan, S. & Parekkadan, B. Non-invasive cell counting of adherent, suspended and encapsulated mammalian cells using optical density. Biotechniques 68, 35–40, doi:10.2144/btn-2019-0052 (2020).

20 Seita, A., Nakaoka, H., Okura, R. & Wakamoto, Y. Intrinsic growth heterogeneity of mouse leukemia cells underlies differential susceptibility to a growth-inhibiting anticancer drug. Plos One 16, e0236534, doi:10.1371/journal.pone.0236534 (2021).

21 Michl, J., Park, K. C. & Swietach, P. Evidence-based guidelines for controlling pH in mammalian live-cell culture systems. Commun Biol 2, 144, doi:10.1038/s42003-019-0393-7 (2019).

22 Rovati, L., Fabbri, P., Ferrari, L. & Pilati, F. Construction and evaluation of a disposable pH sensor based on a large core plastic optical fiber. Rev Sci Instrum 82, 023106, doi:10.1063/1.3541795 (2011).

23 Magnusson, E. B., Halldorsson, S., Fleming, R. M. & Leosson, K. Real-time optical pH measurement in a standard microfluidic cell culture system. Biomed Opt Express 4, 1749–1758, doi:10.1364/BOE.4.001749 (2013).

24 Yus, E. et al. Impact of genome reduction on bacterial metabolism and its regulation. Science 326, 1263–1268, doi:10.1126/science.1177263 (2009).

25 Broto, A., Gaspari, E., Miravet-Verde, S., Dos Santos, V. & Isalan, M. A genetic toolkit and gene switches to limit Mycoplasma growth for biosafety applications. Nat Commun 13, 1910, doi:10.1038/s41467-022-29574-0 (2022).

26 Andersson, L. C., Nilsson, K. & Gahmberg, C. G. K562--a human erythroleukemic cell line. Int J Cancer 23, 143–147, doi:10.1002/ijc.2910230202 (1979).

27 Schneider, U., Schwenk, H. U. & Bornkamm, G. Characterization of EBV-genome negative “null” and “T” cell lines derived from children with acute lymphoblastic leukemia and leukemic transformed non-Hodgkin lymphoma. Int J Cancer 19, 621–626, doi:10.1002/ijc.2910190505 (1977).

28 Rasheed, S., Nelson-Rees, W. A., Toth, E. M., Arnstein, P. & Gardner, M. B. Characterization of a newly derived human sarcoma cell line (HT-1080). Cancer 33, 1027–1033, doi: 10.1002/1097-0142(197404)33:4<1027::aid-cncr2820330419>3.0.co;2-z (1974).

29 Graham, F. L., Smiley, J., Russell, W. C. & Nairn, R. Characteristics of a human cell line transformed by DNA from human adenovirus type 5. J Gen Virol 36, 59–74, doi:10.1099/0022-1317-36-1-59 (1977).

30 P., B. P. a. H. Using Phenol Red to Assess pH in Long-Term Proliferation Assays. BioTek Instruments, Inc., doi:https://www.biotek.com/assets/tech_resources/SLAS%202018%20Phenol%20red-LR.pdf (2018).

31 Zhang, L. et al. Control of IgG glycosylation in CHO cell perfusion cultures by GReBA mathematical model supported by a novel targeted feed, TAFE. Metab Eng 65, 135–145, doi:10.1016/j.ymben.2020.11.004 (2021).

32 Alton, G. et al. Direct utilization of mannose for mammalian glycoprotein biosynthesis. Glycobiology 8, 285–295, doi:10.1093/glycob/8.3.285 (1998).

33 Vergara, M. et al. Differential effect of culture temperature and specific growth rate on CHO cell behavior in chemostat culture. Plos One 9, e93865,doi:10.1371/journal.pone.0093865 (2014).

34 Howard, G. R., Jost, T. A., Yankeelov, T. E. & Brock, A. Quantification of long-term doxorubicin response dynamics in breast cancer cell lines to direct treatment schedules. PLoS Comput Biol 18, e1009104, doi:10.1371/journal.pcbi.1009104 (2022).

35 Tacar, O., Sriamornsak, P. & Dass, C. R. Doxorubicin: an update on anticancer molecular action, toxicity and novel drug delivery systems. J Pharm Pharmacol 65, 157–170, doi:10.1111/j.2042-7158.2012.01567.x (2013).

36 Zwelling, L. A. et al. HT1080/DR4: a P-glycoprotein-negative human fibrosarcoma cell line exhibiting resistance to topoisomerase II-reactive drugs despite the presence of a drug-sensitive topoisomerase II. J Natl Cancer Inst 82, 1553–1561,doi:10.1093/jnci/82.19.1553 (1990).

37 Albright, C. F. et al. Matrix metalloproteinase-activated doxorubicin prodrugs inhibit HT1080 xenograft growth better than doxorubicin with less toxicity. Mol Cancer Ther 4, 751–760, doi:10.1158/1535-7163.MCT-05-0006 (2005).

38 Matreyek, K. A., Stephany, J. J. & Fowler, D. M. A platform for functional assessment of large variant libraries in mammalian cells. Nucleic Acids Res 45, e102, doi:10.1093/nar/gkx183 (2017).

39 Moffat, A. D., Elliston, A., Patron, N. J., Truman, A. W. & Carrasco Lopez, J. A. A biofoundry workflow for the identification of genetic determinants of microbial growth inhibition. Synth Biol (Oxf) 6, ysab004, doi:10.1093/synbio/ysab004 (2021).

40 Dudley, Q. M. et al. Biofoundry-assisted expression and characterization of plant proteins. Synth Biol (Oxf) 6, ysab029, doi:10.1093/synbio/ysab029 (2021).

41 Wong, B. G., Mancuso, C. P., Kiriakov, S., Bashor, C. J. & Khalil, A. S. Precise, automated control of conditions for high-throughput growth of yeast and bacteria with eVOLVER. Nat Biotechnol 36, 614–623, doi:10.1038/nbt.4151 (2018).

42 Fonseca, J. P. et al. A Toolkit for Rapid Modular Construction of Biological Circuits in Mammalian Cells. Acs Synth Biol 8, 2593–2606, doi:10.1021/acssynbio.9b00322 (2019).

43 Martella, A., Matjusaitis, M., Auxillos, J., Pollard, S. M. & Cai, Y. EMMA: An Extensible Mammalian Modular Assembly Toolkit for the Rapid Design and Production of Diverse Expression Vectors. Acs Synth Biol 6, 1380–1392, doi:10.1021/acssynbio.7b00016 (2017).

44 Holowko, M. B., Frow, E. K., Reid, J. C., Rourke, M. & Vickers, C. E. Building a biofoundry. Synth Biol (Oxf) 6, ysaa026, doi:10.1093/synbio/ysaa026 (2021).

45 Le, K. et al. Assuring Clonality on the Beacon Digital Cell Line Development Platform. Biotechnol J 15, e1900247, doi:10.1002/biot.201900247 (2020).

46 Le, K. et al. A novel mammalian cell line development platform utilizing nanofluidics and optoelectro positioning technology. Biotechnol Prog 34, 1438–1446, doi:10.1002/btpr.2690 (2018).

