## Supplementary file for "Mammalian cell characterisation by non-invasive plate reader assay"

### Supplementary Information

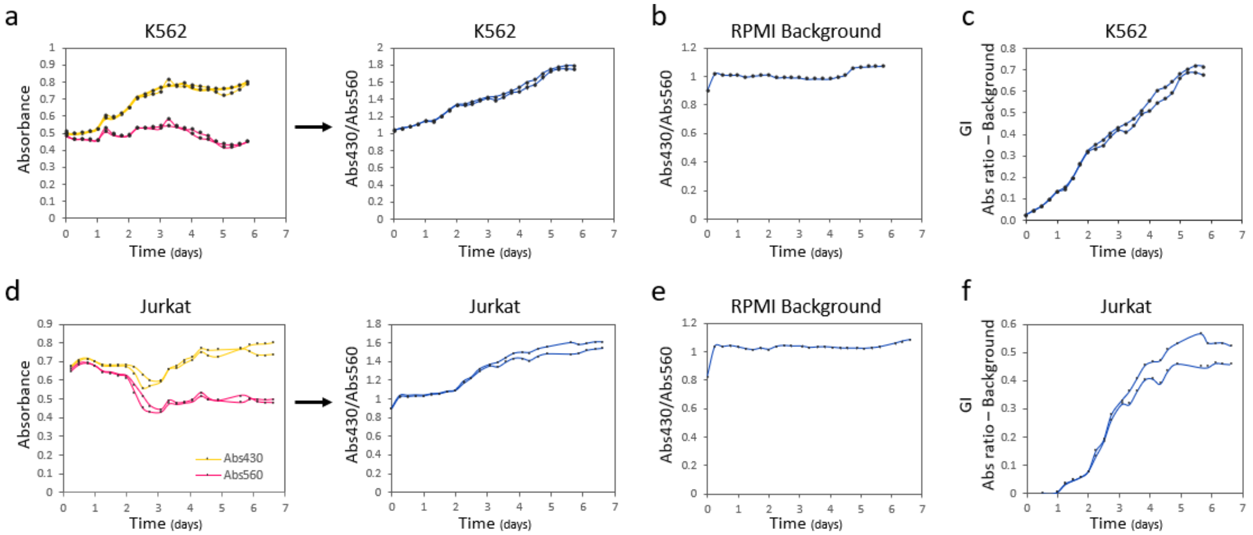

**Figure S1. GI calculation for suspension cells from phenol red individual plate reader absorbance values. a)** Representative duplicate variations of $Abs_{430}$ (yellow) and $Abs_{560}$ (pink) over time (left) for RPMI medium when K562 cells are cultured. Resulting $Abs_{430}$ over $Abs_{560}$ ratios (right, blue) over time. **b)** Average background and error of the mean for $Abs_{430}/Abs_{560}$ ratio of RPMI over time (n=8) in control wells containing no cells. **c)** Representative duplicate GI profiles over time for K562 cells, resulting from $Abs_{430}/Abs_{560}$ ratio normalised to RPMI background. **d)** Representative duplicates of $Abs_{430}$ (yellow) and $Abs_{560}$ (pink) over time (left) for RPMI medium over cultured Jurkat cells. Resulting $Abs_{430}$ over $Abs_{560}$ ratios (right, blue) over time. **e)** Background and error of the mean for $Abs_{430}/Abs_{560}$ ratio of RPMI over time (n=8) in wells containing no cells. **f)** Representative duplicate GI profiles over time for Jurkat cells, resulting from $Abs_{430}/Abs_{560}$ ratio normalised to RPMI background.

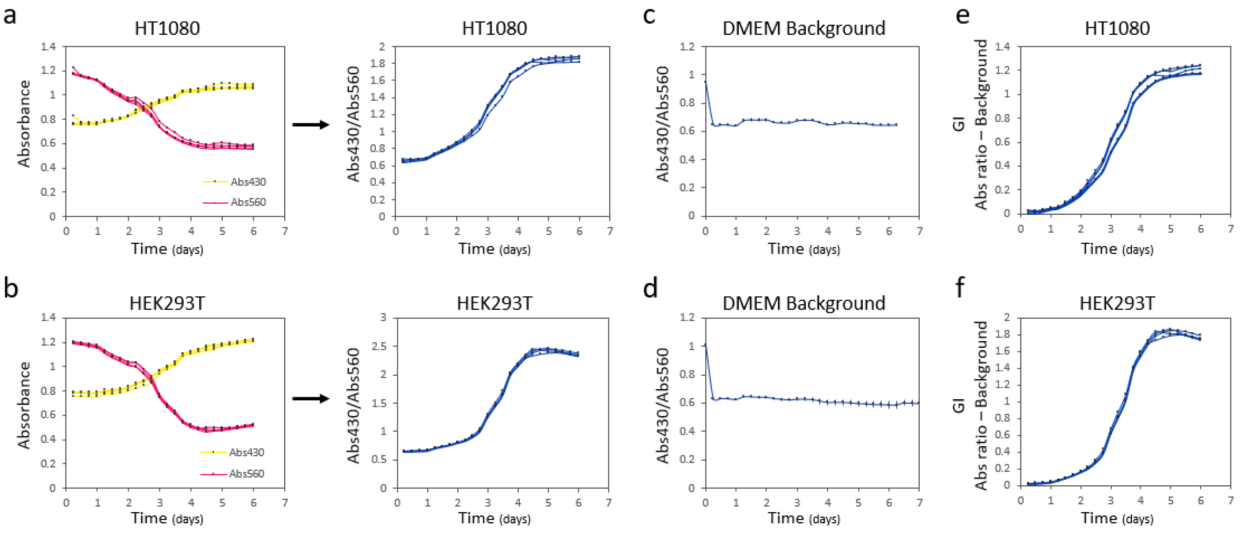

**Figure S2. GI calculation for adhesion cells from phenol red individual plate reader absorbance values.** $Abs_{430}$ (yellow) and $Abs_{560}$ (pink) over time (left) for DMEM medium when HT1080 **(a)** and HEK293T **(b)** cells are cultured. Resulting $Abs_{430}$ over $Abs_{560}$ ratio (right, blue) over time (n=4). Average background and error of the mean for $Abs_{430}/Abs_{560}$ ratio of DMEM over time (n=3) in control wells containing no cells **(c, d)** GI profiles over time for HT1080 **(e)** and HEK293T **(f)** cells, resulting from $Abs_{430}/Abs_{560}$ ratio normalised to DMEM background (n=4).

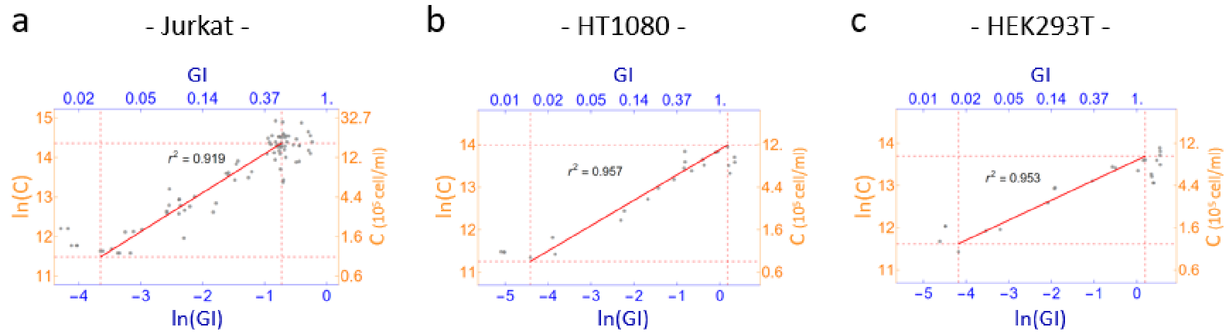

**Figure S3. Linear correlation of GI and C for Jurkat, HT1080 and HEK293T cells grown at 37°C with Glu.** Graphs representing the correlation between GI and C for **a)** Jurkat, **b)** HT1080 and **c)** HEK293T cells in logarithmic scale. Red dashed lines represent the edges of linear region corresponding to the exponential phase. The red solid line is the linear fit of the data in linear region for which $r^2$ values are indicated. Linear fit equations are reported in Table S2. Numbers of biological repeats for each sample are reported in Table S3. Data analysis is described in the Methods section and in Supplementary Note 1.

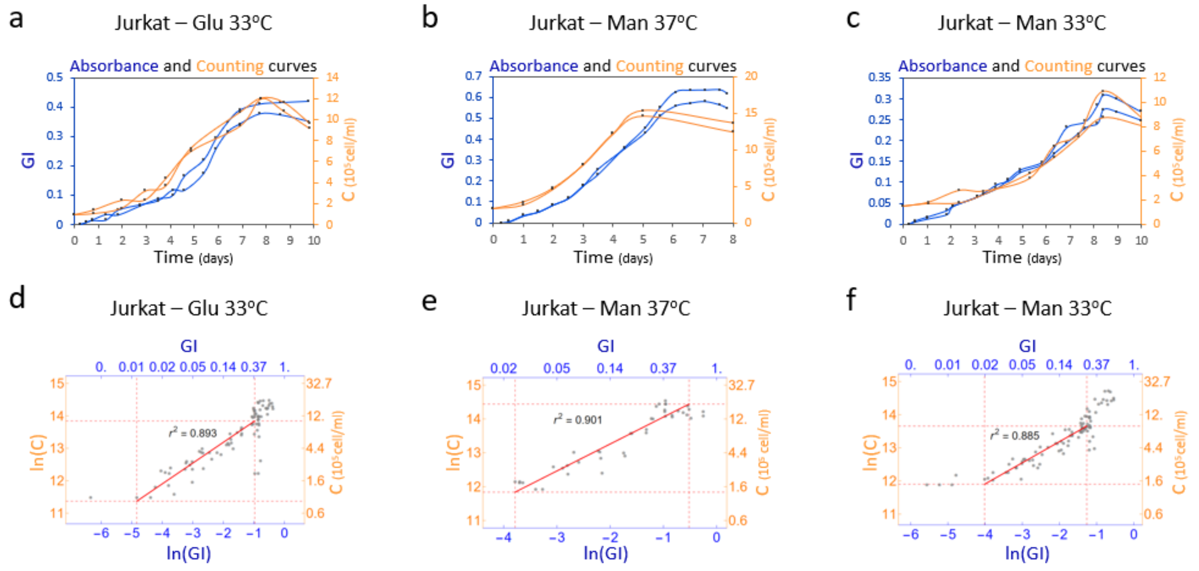

**Figure S4. Growth of Jurkat suspension cells under different conditions can be characterised by a plate reader assay.** Representative GI (blue, left vertical axis) and C (orange, right vertical axis) profiles over time for Jurkat cells grown with Glu at 33°C **(a)** or Man at either 37°C **(b)** or 33°C **(c)**. All biological replicates can be found in Related File 1. Correlation between GI and C for Jurkat cells grown in presence of Glu at 33°C **(d)** or Man at either 37°C **(e)** or 33°C **(f)** in logarithmic scale. Red dashed lines represent the edges of the linear region. The red solid line is the linear fit of the data in the linear region for which $r^2$ values are indicated. Linear fit equations are reported in Table S2. Numbers of biological repeats for each sample are reported in Table S3. Data analysis is described in the Methods section and in Supplementary Note 1.

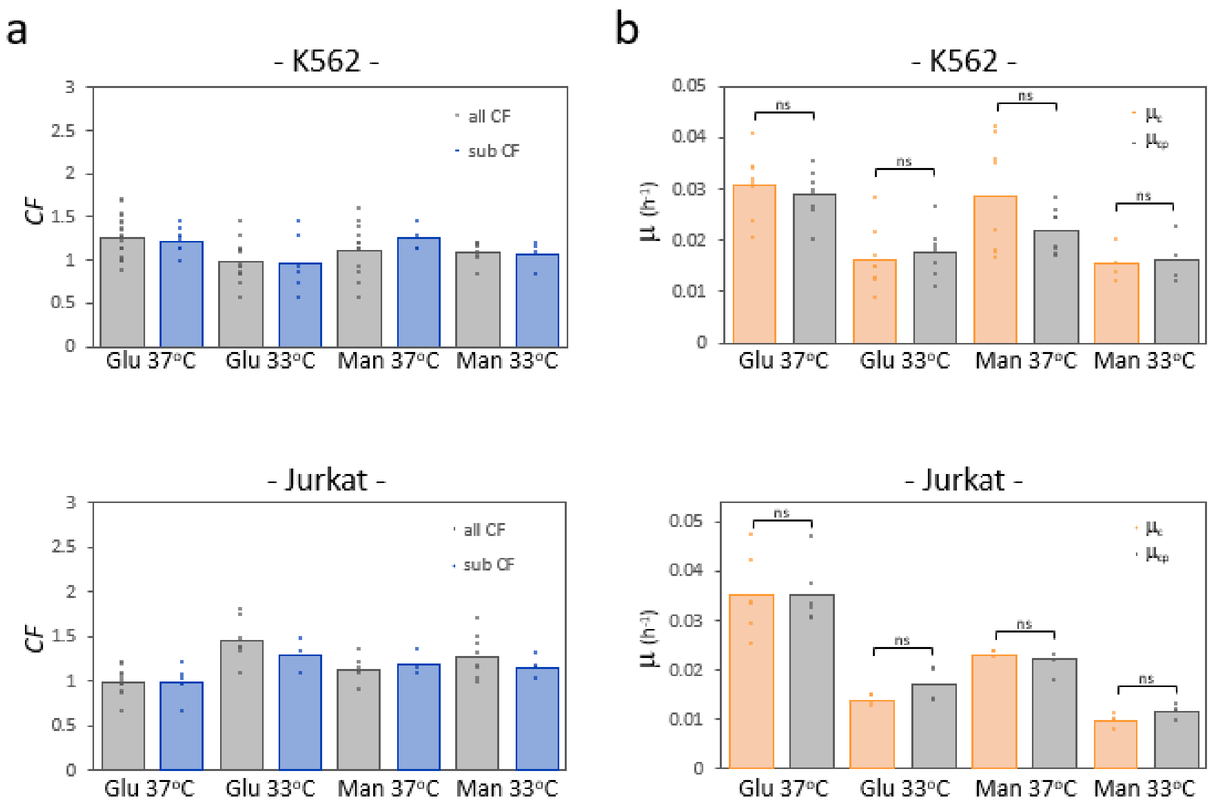

**Figure S5. Reproducibility test of plate reader growth characterisation for K562 and Jurkat cells grown in different conditions.** To test the automatable capability of our plate reader assay, the data set obtained for each growing condition was split. *CF* from one subset were averaged together (Table S5) to convert $\mu_p$ into $\mu_{cp}$ in the remaining data subset. **a)** Bar plots of all (grey) or a subset (sub, blue) of *CF* for K562 (top) and Jurkat (bottom) cells grown in presence of either Glu or Man at 37°C or 33°C. **b)** Bar plots of $\mu_c$ (orange) and $\mu_{cp}$ (grey), calculated as $\mu_p$ x CF average, for K562 (top) and Jurkat (bottom) cells grown in presence of either Glu or Man at 37°C or 33°C are shown. A two-sided t-test indicated that average *CF* of all or one subset are compatible ($p>0.05$) and that $\mu_{cp}$ are compatible with actual $\mu_c$ (ns, non-significant: $p>0.05$). The height of the bars represents the average value of the single replicates shown as dots. Numbers of biological repeats for each sample are reported in Table S3. Data analysis is described in the methods section and in Supplementary Note 1.

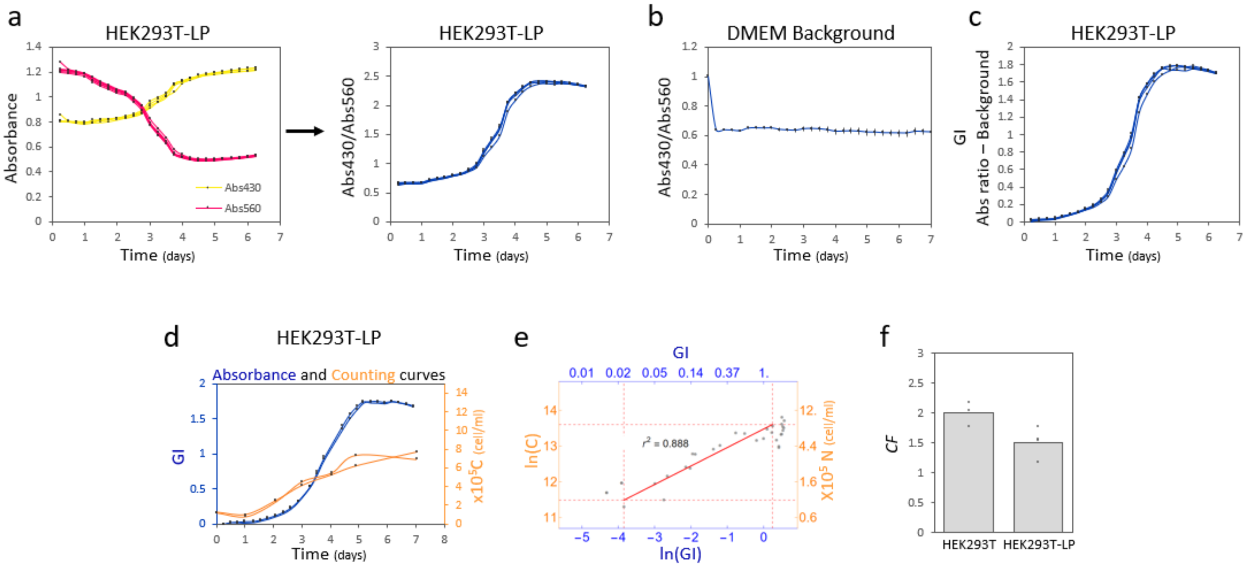

**Figure S6. Growth of HEK293T-LP cells can be characterised with the plate reader-based assay. a)** $Abs_{430}$ (yellow) and $Abs_{560}$ (pink) over time (left) for DMEM medium when HEK293T-LP cells are cultured. Resulting $Abs_{430}$ over $Abs_{560}$ ratio (right, blue) over time (n=4). **b)** Average background and error of the mean for $Abs_{430}/Abs_{560}$ ratio of DMEM over time (n=3) in control wells containing no cells.. **c)** GI profiles over time for HEK293T-LP cells, resulting from $Abs_{430}/Abs_{560}$ ratio normalised to DMEM background (n=4). **d)** Representative growth curves resulting from phenol red acidification (GI, blue) and cell counts (C, orange). All biological replicates can be found in Related File 1. **e)** Correlation between GI and C in logarithmic scale. Red dashed lines represent the edges of the linear region. Red solid line is the linear fit of the data in linear region for which $r^2$=0.888. The linear fit equations are reported in Table S2. **f)** Bar plot of *CF* for HEK293T-LP cells compared to their HEK293T parental cell line. The height of the bar represents the average value of the single replicates shown as black dots. Numbers of biological repeats for each sample are reported in Table S3. Data analysis is described in the Methods section and in Supplementary Note 1.

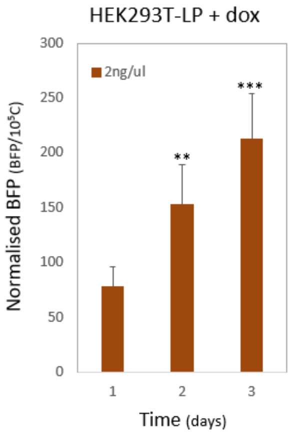

**Figure S7. BFP expression level in HEK293T-LP cells after dox induction.** Normalised fluorescence per cell (calculated as BFP/$10^5$C) for HEK293T-LP cells induced with 2ng/µl dox over three days. A two-sided t-test indicated that normalised BFP levels increase significantly each day (**: $p<0.01$ and ***: $p<0.001$). Numbers of biological repeats for each sample are reported in Table S3. Data and statistical analysis are described in the Methods section.

### Supplementary tables

| Cell line and growing condition | GI interval of linearity $[GI_{min}; GI_{max}]$ | C interval of linearity $[C_{min}; C_{max}]$ ($10^5$ cell/ml) |
|---|---|---|
| K562 Glu37°C | [0.034; 0.48] | [1.02; 10.7] |
| Jurkat Glu 37°C | [0.026; 0.48] | [0.97; 17.3] |
| HT1080 Glu 37°C | [0.012; 1.19] | [0.77; 11.9] |
| HEK293T Glu 37°C | [0.015; 1.21] | [1.1; 8 .8] |
| HEK293T-LP Glu 37°C | [0.021; 1.28] | [0.97; 8.1] |
| K562 Glu 33°C | [0.022; 0.340] | [0.75; 11.5] |
| Jurkat Glu 33°C | [0.008; 0.378] | [0.87; 10.3] |
| K562 Man 37°C | [0.035; 0.306] | [1.2; 11.2] |
| Jurkat Man 37°C | [0.023; 0.596] | [1.4; 18.7] |
| K562 Man 33°C | [0.019; 0.265] | [0.98; 10.8] |
| Jurkat Man 33°C | [0.018; 0.283] | [1.5; 8.5] |

**TABLE S1.** Intervals of linearity for each condition: Minimum and maximum values of GI and C in which ln(C) vs ln(GI) show the best linear trend (highest $r^2$ which values are reported in **TableS2**).

| Cell line and growing condition | Fit ln(C) = a + b*ln(GI) | $r^2$ | Fit parameter (a ± erra) | Fit parameter (b ± errb) |
|---|---|---|---|---|
| K562 Glu 37°C | ln(C) = 14.5 + 0.88*ln(GI) | 0.915 | (14.5 ± 0.1) | (0.88 ± 0.03) |
| Jurkat Glu 37°C | ln(C) = 15.1 + 0.99*ln(GI) | 0.919 | (15.1 ± 0.1) | (0.99 ± 0.04) |
| HT1080 Glu 37°C | ln(C) = 13.9 + 0.60*ln(GI) | 0.957 | (13.9 ± 0.1) | (0.60 ± 0.03) |
| HEK293T Glu 37°C | ln(C) = 13.6 + 0.47*ln(GI) | 0.953 | (13.6 ± 0.1) | (0.47 ± 0.03) |
| HEK293T-LP Glu 37°C | ln(C) = 13.5 + 0.52*ln(GI) | 0.888 | (13.5 ± 0.1) | (0.52 ± 0.05) |
| K562 Glu 33°C | ln(C) = 15.0 + 0.99*ln(GI) | 0.861 | (15.0 ± 0.1) | (0.99 ± 0.04) |
| Jurkat Glu 33°C | ln(C) = 14.5 + 0.64*ln(GI) | 0.893 | (14.5± 0.1) | (0.64 ± 0.03) |
| K562 Man 37°C | ln(C) = 15.2 + 1.03*ln(GI) | 0.909 | (15.2 ± 0.1) | (1.03 ± 0.05) |
| Jurkat Man 37°C | ln(C) = 14.9 + 0.80 * ln(GI) | 0.901 | (14.9 ± 0.1) | (0.80 ± 0.04) |
| K562 Man 33°C | ln(C) = 15.1 + 0.91*ln(GI) | 0.928 | (15.1 ± 0.1) | (0.91 ± 0.03) |
| Jurkat Man 33°C | ln(C) = 14.5 + 0.64*ln(GI) | 0.885 | (14.5 ± 0.1) | (0.64 ± 0.03) |

**TABLE S2**. Results of the best fits for determining the relation between ln(C) and ln(GI) for all growing conditions.

| | Number of biological repeats | |
|---|---|---|
| **Cell line and growing condition** | Plate reader | Cell counting |
| K562 Glu 37°C | 20 | 20 |
| Jurkat Glu 37°C | 12 | 12 |
| HT1080 Glu 37°C | 3 | 3 |
| HEK293T Glu 37°C | 3 | 3 |
| HEK293T-LP Glu 37°C | 4 | 4 |
| K562 Glu 33°C | 11 | 11 |
| Jurkat Glu 33°C | 7 | 7 |
| K562 Man 37°C | 12 | 12 |
| Jurkat Man 37°C | 6 | 6 |
| K562 Man 33°C | 8 | 8 |
| Jurkat Man 33°C | 8 | 8 |
| HT1080 + doxo (0, 10, 25, 50, 75, 100, 250, 500, 750 and 1000nM) | 4 | - |
| K562 + doxo (0, 10, 25, 50, 75, 100, 250, 500, 750 and 1000nM) | 4 | - |
| HEK293T-LP + dox (0, 0.1, 0.25, 0.5, 1 and 2ng/ul) | 7 | - |

**Tables S3.** Number of biological repeats per experiments with plate reader and cell counting protocol summary.

| | Average ± err |
|---|---|
| K562 Glu 37°C | (1.26 ± 0.05) |
| Jurkat Glu 37°C | (0.98 ± 0.04) |
| HT1080 Glu 37°C | (1.78 ± 0.08) |
| HEK293T Glu 37°C | (2.00 ± 0.12) |
| HEK293T-LP Glu 37°C | (1.51 ± 0.12) |

**Tables S4.** Average *CF* for cells grown at 37°C with Glu.

| | Average of all *CF* | Average of *CF* subset |
|---|---|---|
| K562 Glu 37°C | (1.26 ± 0.05) | (1.22 ± 0.05) |
| K562 Glu 33°C | (0.97 ± 0.04) | (1 ± 0.04) |
| K562 Man 37°C | (1.12 ± 0.09) | (1.05 ± 0.12) |
| K562 Man 33°C | (1.09 ± 0.04) | (1.11 ± 0.04) |
| Jurkat Glu 37°C | (0.98 ± 0.04) | (0.99 ± 0.05) |
| Jurkat Glu 33°C | (1.45 ± 0.09) | (1.57 ± 0.12) |
| Jurkat Man 37°C | (1.13 ± 0.06) | (1.07 ± 0.09) |
| Jurkat Man 33°C | (1.27 ± 0.09) | (1.39 ± 0.15) |

**Tables S5.** Average *CF* values across the different growing conditions of K562 and Jurkat cells.

### Supplementary Note 1. Computational pipeline for automated data analysis

**Characterisation of the relation between GI and C.** In Figure S8, we show an example of the procedure used to obtain the final relation between ln(GI) and ln(C). It is applied to the data set of K562 cells growing in Glu at 37°C.The procedure consists of two main steps: i) determination of the best region of linearity between the two variables and ii) linear fit of the data belonging to such region in order to quantify the relation between ln(GI) and ln(C).

**i) Identification of the best range of linearity.** Our method is based on the automatic identification, through an iterative process, of the subset of data whose linear fit gives the highest coefficient of determination, $r^2$. To do this, we first excluded outliers with GI < $GI_0$ or C < $C_0$, where $C_0$ is the initial concentration of cells and $GI_0$ is GI normalised to zero at six hours. The constraint on C was relaxed in the analysis of adherent cell lines to account for a smaller dataset with respect to suspension cell lines. Then, we identified the two data with the lowest ($d_{min}$) and highest ($d_{max}$) value of ln(GI) (Figure S8a). By looping from the lowest to the highest ln(GI) value, a linear fit of a subset of the data was performed and its $r^2$ was computed. The loop started by considering at least the first n= 5 sorted data-points (i.e. those with the five lowest values of ln(GI) coordinate) to minimize $r^2$ bias. At each step of the loop, the set of data was increased of one element by increasing the ln(GI) coordinate. In this way, the trend of $r^2$ as a function of the maximum ln(GI) value of each fitted data-subset was obtained (Figure S8b). The value of ln(GI) corresponding to the fit with the highest $r^2$, was considered as the upper bound of the linear fit ($d_u$, red point and red vertical line in Figure S8b). Analogously, we determined the lower bound of the linear region. In this case, only data with ln(GI) ≤ $d_u$ were considered and linear fits were performed by looping on the data sorted from the smallest ln(GI) to ln(GI) = $d_u$. A loop was performed by, at each step, i) increasing the value of ln(GI), ii) consider the data with ln(GI) bigger than the value set at point i), and iii) perform a linear fit and computing its $r^2$ value. The loop was stopped when the dataset to be fitted had n elements. In this way, the trend of $r^2$ as a function of ln(GI) was obtained (Figure S8c) and the ln(GI) corresponding to the point with the biggest $r^2$ was considered as the lower bound of the linearity region ($d_l$, red point and its correspondent red vertical line in Figure S8c). The resulting intervals of linearity for each of the studied conditions are reported in Table S1, while the values of $r^2$ are reported in Table S2.

**ii) Relation between C and GI.** Once the edges of the region of linearity between ln(GI) and ln(C) were determined, we fitted the data with a final linear fit that sets the quantitative relation between the two variables (Figure S8d), of the form:

$$ln(C) = a + b*ln(GI) \qquad (1)$$

The values of the fitted parameters *a* and *b* with their errors for all the studied conditions are listed in Table S2.
From (1) it follows:
$C = \exp(a)*GI^b$. (2)

**Exponential growth rates and *CF*.**
As shown by Figure S8e, the range of linearity corresponds to the exponential phase of growth, before saturation. Thanks to this, it is possible to map $\mu_p$ into $\mu_c$ through a simple proportion. Since a linear relation allows to map ln(GI) into ln(C), any linear relation found in the space ln(GI) *vs* time can be linearly mapped into the space ln(C) *vs* time.
The growth curves used to compute growth rates were expressed in semi-logarithmic scale and the data were normalised by their initial value ($C_0$ or $GI_0$). Indeed, the growth curve of cell counts was expressed as $\ln(C/C_0)$ *vs* time and that of plate reader was $\ln(GI/GI_0)$ *vs* time, where $C_0$ and $GI_0$ are values of C at zero hour and GI at six hours respectively (Related file 1). For plate reader measurements, we discarded the value at zero hour since it corresponded to an outlier value.
To obtain the exponential growth rate, we fitted with a line the data of ln(C) and ln(GI) growth curves belonging to the exponential phase of growth. An automatic and analytic procedure to determine the exponential phase of growth is the one we previously adopted in *Enrico Bena et al*[1], where the growth curves were fitted by using a logistic function whose parameters were the lag time, the exponential growth rate and the level of saturation. The edges of the exponential phase of growth were analytically determined from the logistic function. Growth curves resulting from cell counts rarely showed a lag-phase and saturated within three time points. Given this, we couldn't use the analytic procedure to infer the edges of the exponential phase, so we set it manually per each growth curve, paying attention to include in the exponential phase of growth at least three data points. The slope of the linear fit of the cell counts data within the exponential phase of growth is $\mu_c$. Then, we selected the data of plate reader growth curves that had a correspondent value in cell counts and belonged to the exponential phase of growth and fitted them with a linear fit to obtain $\mu_p$. The CF that converts $\mu_p$ into $\mu_c$ were then computed as the ratio: $CF = \mu_p/\mu_c$.
Concerning analysis of cells treated with doxo, in order to obtain the values shown in Figure 3b, we i) first computed all the growth rates with the analytic procedure mentioned above, then ii), per each run, we computed the average growth rate of cells grown with 0 nM of doxo and iii) we normalized all the growth rates of the same run to this average value and expressed it in percentage.

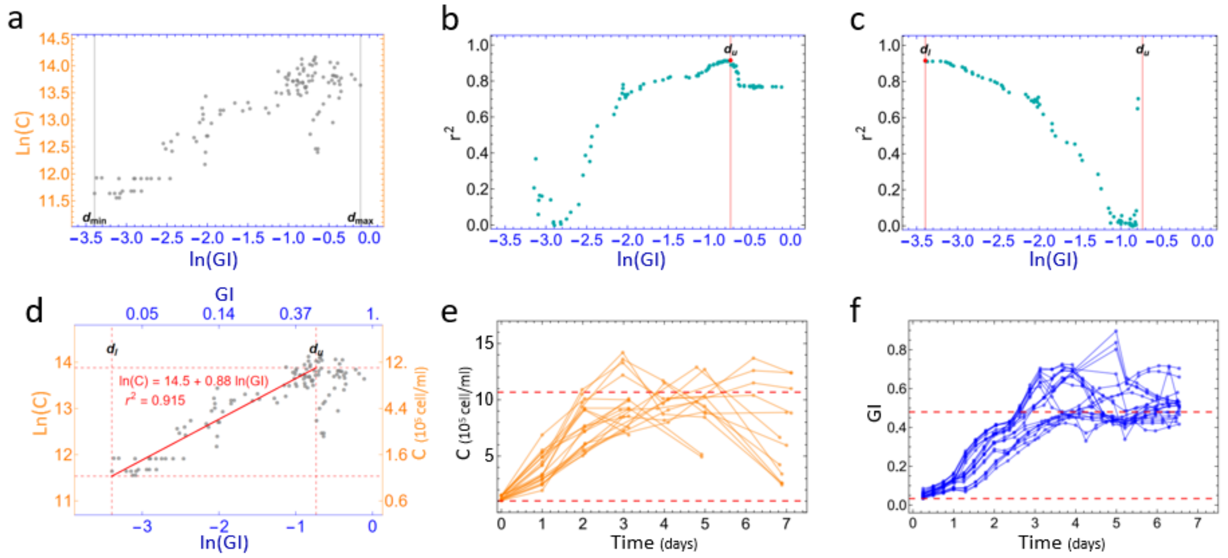

**Figure S8**. **Steps to characterise the relation between GI and C.** Data shown are K562 cells grown in Glu at 37°C (20 independent growth curves belonging to 5 different runs). **a)** ln(C) is plotted as a function of the ln(GI). The edges of the data in ln(GI) are selected ($d_{min}$ and $d_{max}$) and emphasized by vertical grey lines. **b)** $r^2$ plotted as a function of the biggest value of the subset of fitted data. $d_u$ is emphasized in red and sets the upper bound of the linear region (vertical red line). **c)** Analogously to **b)**, the highest $r^2$ sets $d_l$. $d_l$ is emphasized in red and by the red vertical line. **d)** Linear fit of the data within the linear region. Dashed red lines set the edges of the region determined by $d_u$ and $d_l$. The equation of the final and best linear fit is expressed in the figure as well as the $r^2$ value. The limits of the linear region are emphasized as horizontal dashed red lines on cell counts **(e)** and plate reader **(f)** growth curves.

### Additional manuscript Related files

**Related file 1. Plots of ln(GI) and ln(C) over time for all biological replicates.** Available upon requests to authors.

**Related file 2. Excel file of $\mu_p$, $\mu_c$ and *CF* for all biological replicates.** Available upon requests to authors.

**Related file 3. *Wolfram Mathematica* code used for data analysis.** Available upon requests to authors.

### Supplementary References

1 Enrico Bena, C. *et al.* Initial cell density encodes proliferative potential in cancer cell populations. *Sci Rep* **11**, 6101, doi:10.1038/s41598-021-85406-z (2021).
